# Differential Activity of Repurposed Drugs as Receptor Binding Domain Antagonists for *Omicron* and Native Strains of SarsCov2

**DOI:** 10.1101/2022.03.09.483630

**Authors:** Kranti Meher, Saranya K, Arpitha Reddy, Gopi Kadiyala, Subra Iyer, Subhramanyam Vangala, Satish Chandran, Uday Saxena

## Abstract

Omicron strain is the latest variant of concern of SarsCov2 virus. The mutations in this strain in the S protein Receptor Binding domain (RBD) enable it to be more transmissible as well as escape neutralizing activity by antibodies in response to vaccine. Thus, Omicron specific strategies are need to counter infection by this strain.

We investigated a collection of approved drugs shown to antagonize the binding of native strain RBD to human ACE2, for their ability to antagonize binding to Omicron strain RBD.

While most of the drugs the drugs that antagonize binding to native RBD are also active for Omicron RBD but some were inactive, namely drugs that contain iodine are completely inactive against Omicron RBD. Our data strongly indicate that presence of a single iodine molecule in the drug renders it inactive against Omicron strain. Thus, there is molecular specificity of drugs for antagonizing Omicron strain RBD versus native strain RBD of this virus. Such information will pave way for specific drugs for Omicron. A pragmatic message from our data is that the often-used iodine containing mouth wash and rises may be ineffective in antagonizing receptor binding of Omicron strain.

## Introduction

Vaccines against the spike protein of native viruses have become the mainstay of immunization strategy. But recent emergence of newer strains such as the delta and Omicron strain has shown that these antibodies generated against these vaccines have reduced neutralizing activity for the strain, thus leaving open the possibility of fresh round of infections (1). The reason for reduced immune response to newer strain of these vaccines could be the 30 plus mutations that occur in omicron spike protein (2). Specifically, there are 15 amino acid changes in the omicron RBD regions compared to the native RBD. These amino acid changes confer higher affinity binding of its spike protein to human ACE2 receptor and higher transmissibility. Currently 80% of the world’s SarsCov2 virus presence is in the form of omicron strain.

The big challenge with omicron strain is whether we will need new vaccines to fully combat it. It’s possible that reduced immune response that is being seen to this strain may suggest the requirement of strain specific vaccines for full protection. Obviously, this is both demanding and time consuming. Another approach to antagonize new strains is to find drugs that could block the binding of omicron strain RBD to ACE2 receptor and thus prevent viral entry into host cells. This approach can be faster than developing a new vaccine especially if drugs that are currently approved for other indications could be repurposed for antagonizing omicron strain RBD interaction with ACE2.

The effectiveness of vaccines depends heavily on the recipient’s immune system which may be less efficient in pediatric population, elderly and those who are immuno compromised. Therefore, especially in such situations, it would be ideal to have small molecule drugs that could be used to block the entry of the virus both by nasal route or as oral drugs that could act in the oropharyngeal space.

In the studies presented here we tested a collection of available drugs for their ability to antagonize the binding of native strain RBD and Omicron strain RBD to human ACE2. Remarkably we find that while most of the drugs were active for both strains, a collection of chemically diverse drugs which contain iodine were completely inactive in antagonizing Omicron strain RBD binding. Our data shows that presence of iodine molecules in the drug structure inactivates Omicron antagonism while retaining the activity for native RBD. This is first demonstration of a role for iodine in antagonist activity against Omicron. We propose that there are subtle structural and molecular differences for drugs to antagonize the binding of native and Omicron RBD to human ACE2.

## Methods

The antagonist activity of drugs against the binding RBD to human ACE2 was measured using a dot blot assay as indicated before (3). Briefly in the first step, a mixture of horse radish peroxidase-RBD (HRP-RBD) and drugs (drugs at the concentrations of 100μM) were incubated at 37°C for an hour to allow the binding of drugs to HRP-RBD. Following the incubation, these mixtures were added to a nitrocellulose membrane, coated with human ACE-2 receptor to permit the binding of any free HRP-RBD. After incubating the microplate at 37°C for an hour, the membrane was washed four times using wash buffer in order to remove any unbound

HRP-RBD drug complexes. Washing step was followed by addition of a colour substrate; tetramethyleneblue (TMB), which yields blue colour. The reaction was allowed to run for 15 minutes for colour development. This final solution was read at absorbance 450 nm. The colour intensity which is quantitated by ImageJ analysis, is inversely proportional to the inhibition of RBD’s binding to ACE2 by the drug. Data is presented as percent inhibition relative to RBD added without the drug.

The ELISA assay procedure used in the studies has been reported before (3,4).

## Results

### 1. Binding of ACE2 to different Omicron RBD concentrations

To assess the antagonism activity of drugs we used a dot blot assay. RBD was spotted on nitrocellulose membrane and HRP labeled ACE2 was added to measure binding. As shown in Figure 1 the binding of HRP labelled ACE2 to increasing concentrations of Omicron RBD was linear and started to plateau at higher concentrations suggestive of a specific and saturable binding. For the rest of the experiments, we routinely used 0.25 ug/ml of RBD.

**Figure1:**
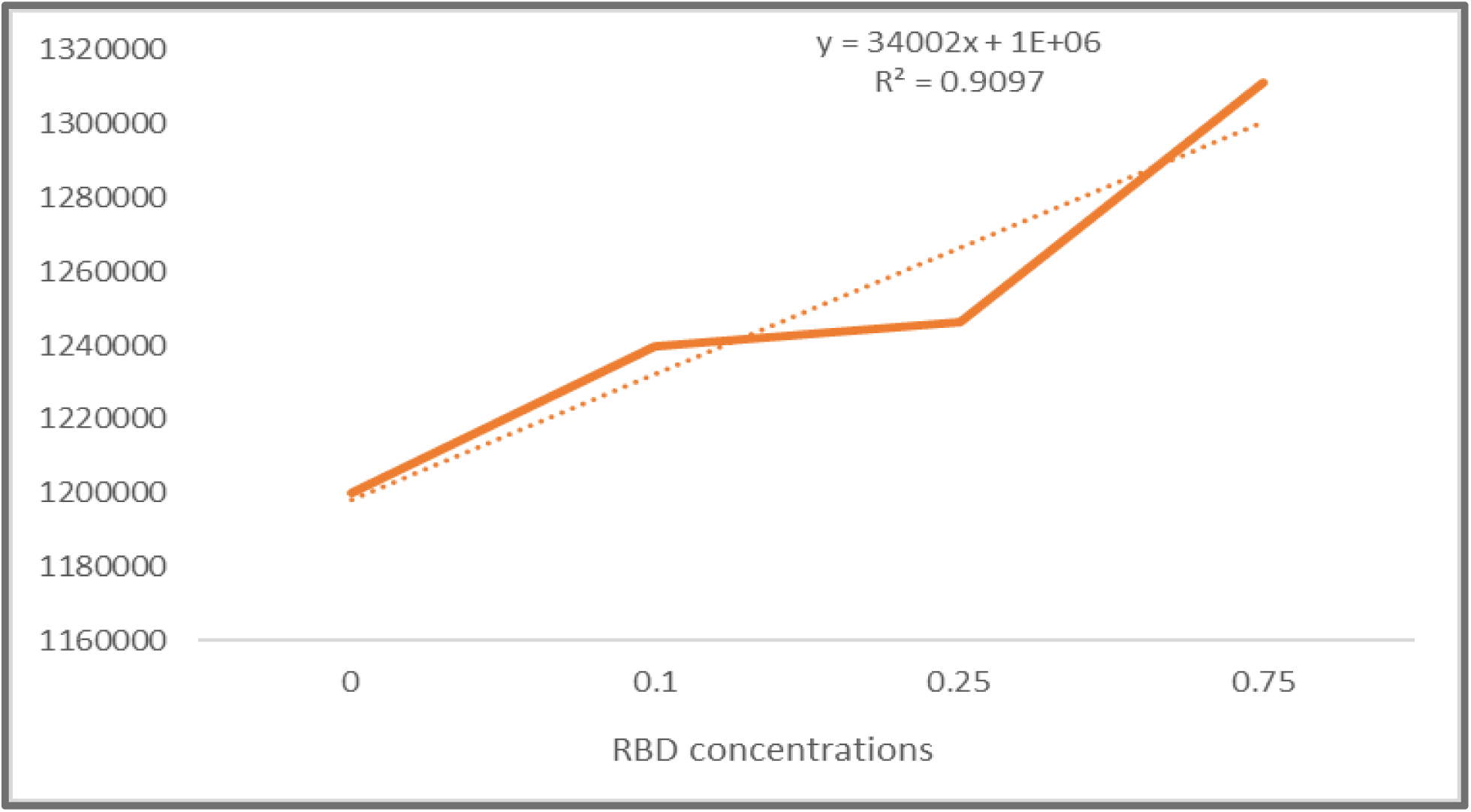
Binding curve of ACE2 (0.25 ug/mL) to different Omicron RBD concentrations in ug/mL.

### 2. Antagonism of RBD binding to ACE2 by drugs

We then tested the antagonist activity of various drugs on the binding of ACE2 to either Omicron RBD or Native RBD. Figure 2 shows that of the

**Figure 2:**
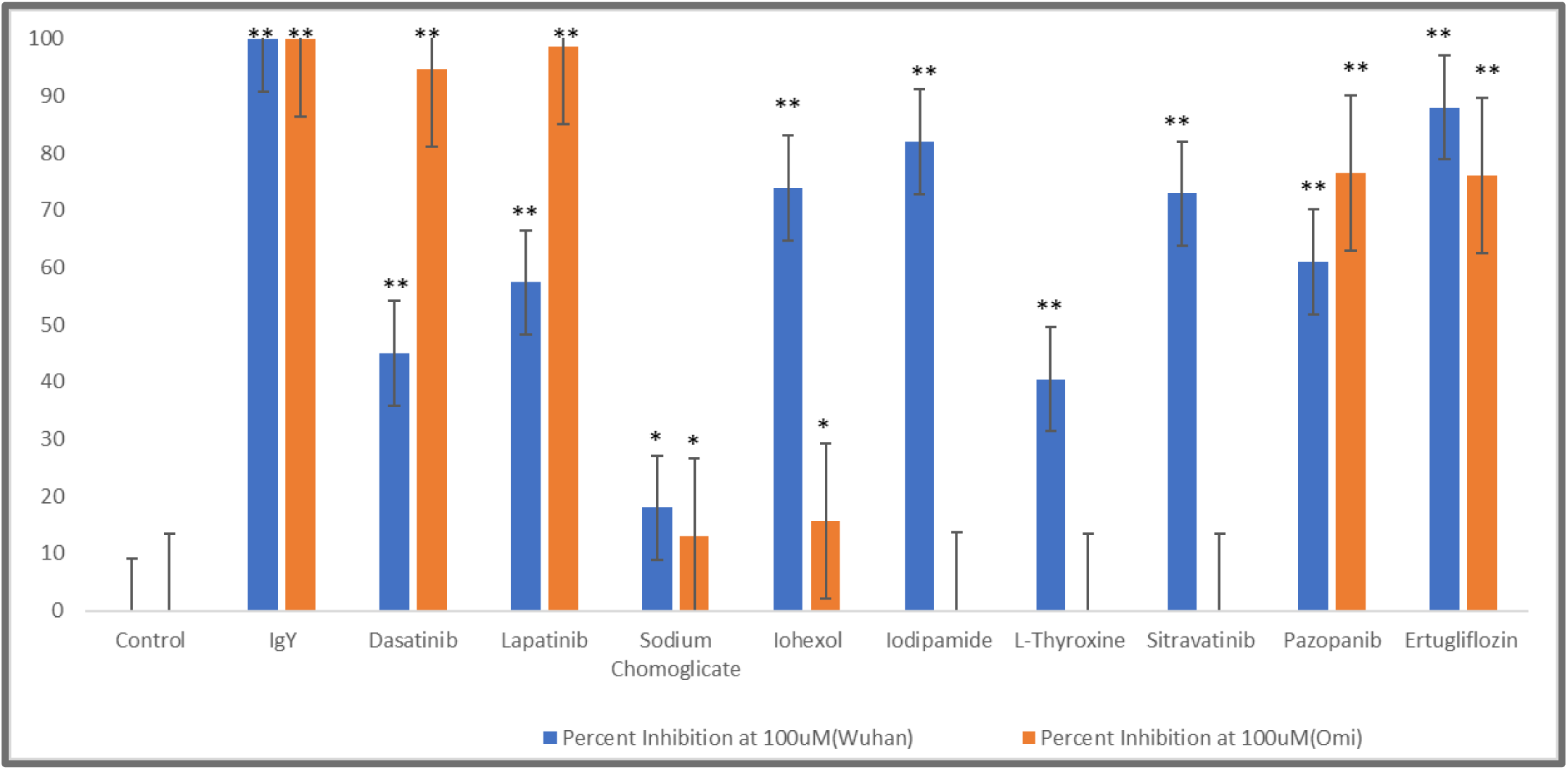
Antagonism of RBD binding (0.25 ug/mL) to ACE2 (0.25 ug/mL) by drugs (100 uM). *** P<0.001; ** P<0.01; *P<0.05. Student’s t-tests were performed to compare between the control (ACE-2+ RBD) and different drugs.

### 3. Comparison Table of drug antagonism activity for Omicron versus native RBD

Table 1 below shows the comparative quantitative activity of the drugs on Omicron versus Native RBD.

**Table 1.**

| **Drug Name** | **Percent Inhibition at 100 uM (Native)** | **Percent Inhibition at 100 uM (Omicron)** |
| --- | --- | --- |
| Dasatinib | 45 | 95 |
| Lapatinib | 57 | 99 |
| Sodium Cromoglicate | 18 | 13 |
| Iohexol | 74 | 16 |
| Iodipamide | 82 | 0 |
| L-Thyroxine | 41 | 0 |
| Sitravatinib | 73 | 0 |
| Pazopanib | 61 | 77 |
| Ertugliflozin | 88 | 76 |

It is evident from the quantitation of drug antagonism summary Table 1 that

1. Dasatinib, Lapatinib, Sodium Cromoglicate, Pazopanib and Ertugliflozin which are structurally diverse all some antagonistic activity ranging from moderate to robust against both Native and Omicron strain RBD. It also underlines the fact that there are no common structural chemical motifs important in the activity
2. Iohexol, Iodipamide, L-Thyroxine and Sitravatinib in contrast are active against Native RBD but drastically reduced or have NO activity against Omicron strain RBD

### 4. Structural similarity of the drugs that are ineffective in Omicron binding

To gain better understanding of any structural commonality in the inactive drugs, we examined the structures of drugs that were active with native RBD but inactive with Omicron RBD. Figure 3 below shows the chemical structures of all 4 inactive drugs

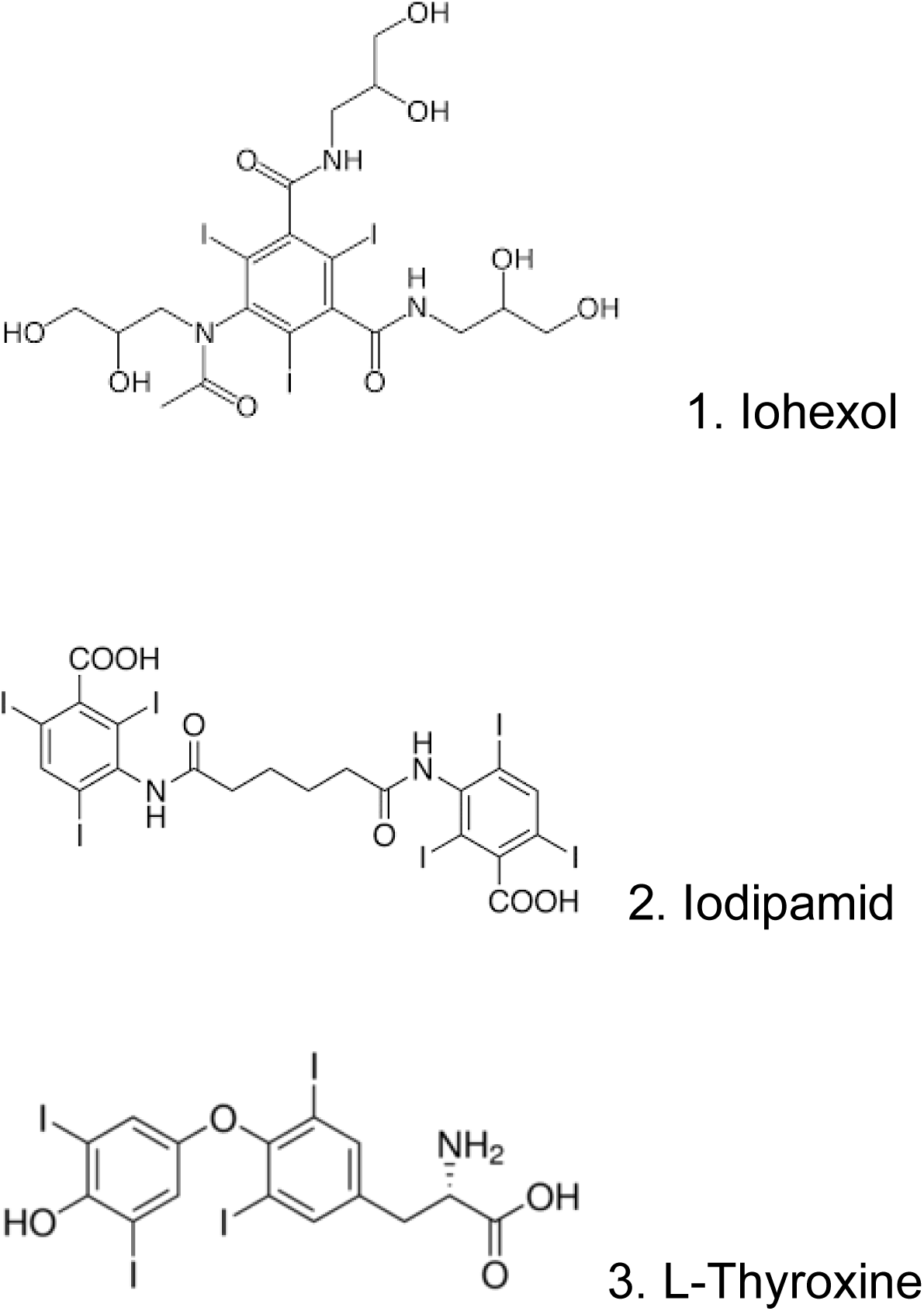
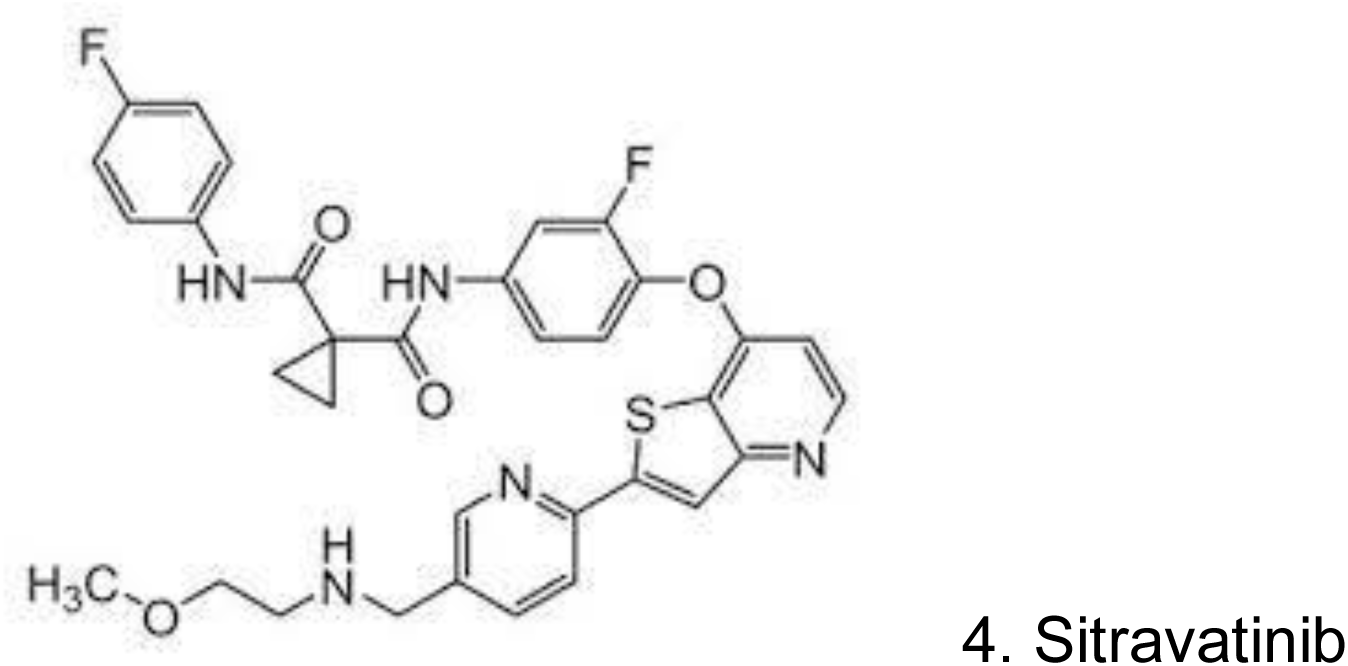

The first thing to note is that all of the structures are very diverse with no obvious relationship to each other. But what is a common feature among all four inactive drugs is the presence of multiple halogen molecules, in case of Iohexol, Iodipamid, L-Thyroxine its iodine molecules and for Sitravatinib it is fluoride. Thus, our data suggest that the presence of Iodine/Fluorine in the drug renders them inactive for Omicron but maintains activity for native strain RBDs.

### 5. Exploration of drug inhibitory activity using an ELISA system

To further explore the nature of interaction between RBD antagonism by the drugs we explored if the drugs can block the binding of RBD to an RBD specific polyclonal IgY antibody (4). This IgY has been previously shown by us to neutralize the binding of both Native and Omicron RBD to ACE2 very effectively.

As shown in Figure 4 below, none of the drugs were significantly different in their ability block the binding of IgY to Native or Omicron RBD. This is unlike drug antagonism of RBD to ACE2 (Table 1) where the drugs were active in Native RBD but inactive in Omicron RBD studies. Thus, there is a clear divergence in the ability of drugs ability to block RBD binding to ACE2 versus RBD binding to ACE2. This suggests that the molecular interaction of the drugs with RBD is to a very narrow region of the protein critical in binding to ACE2. In case of IgY, the binding is likely mediated by multiple epitopes (regions) and therefore not amenable to disruption by the drugs. The take home message from this data is that drugs mediate antagonism action of RBD-ACE2 binding thru on a narrow portion of RBD.

**Figure 4:**
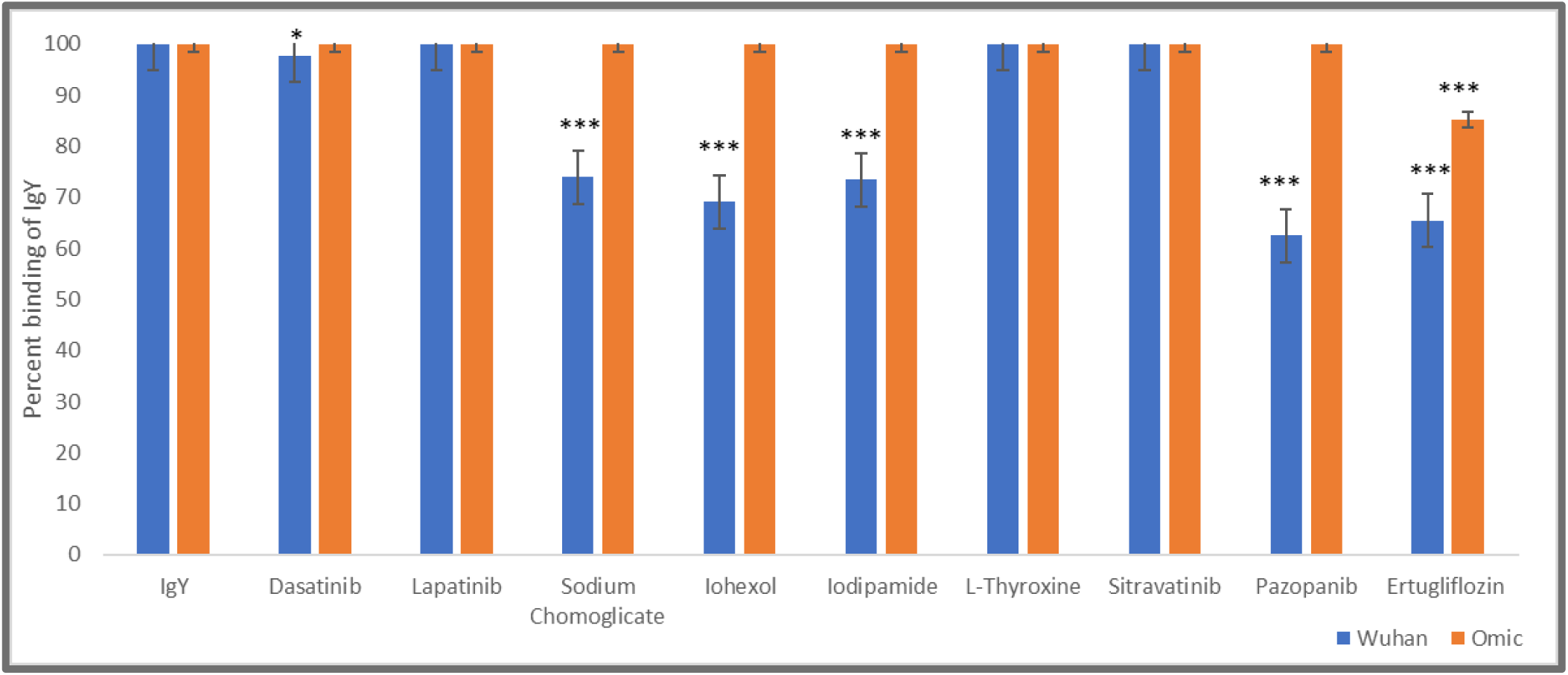
ELISA. Binding of igy to native or omicron rbd and disruption by drugs. *** p<0.001; ** p<0.01; *p<0.05. Student’s t-tests were performed to compare between the control (igy at 2 ug/ml) and different drugs.

## Discussion

Our main objective was to study if selected approved drugs could be repurposed in antagonizing the binding of Omicron strain RBD to human ACE2. As part of this work we compared the inhibitory activity of selected drugs on the binding of native versus Omicron strain RBD binding to human ACE2 receptor. While majority of the drugs were similarly active for both strains, what stood out was the complete lack of activity of iodine containing drugs against Omicron relative to good activity against native strain. This suggests a sophisticated level of differentiation between drug antagonism requirements for Omicron versus native.

One potential reason for this inactivity of iodine containing drugs against Omicron RBD is that iodine as an ion carries a negative charge and this could be the reason for inactivity of iodine containing drugs. But we tend to believe that the negative charge is not the reason for inactivity since other negatively charged drugs are active.

A more intriguing hypothesis for loss of activity for Omicron is that as part of the mutations in the RBD region of Omicron, there are critical tyrosine residues which are important in stabilizing the interactions between RBD and ACE2 (2). Iodine is known to interact with tyrosine residues and perhaps iodine is engaged with tyrosine residues in RBD and ACE2 and therefore the drug is unable to exert antagonistic activity. This remains to be proven in future studies.

Interestingly it has been documented that hyper and hypothyroid conditions in which the iodine containing hormone thyroxine levels fluctuate has been implicated in COVID19 (5). Whether the hormone thyroxine has any interaction with the virus in the body remains to be established.

Another extension of our results pertains to the use of iodine containing oral wash and rinse as a way of blocking the virus’s entry from the buccal cavity. Our data would suggest that at least for omicron strain, iodine containing oral rinse may not be effective.

The inability of drugs to disrupt the binding of IgY to RBD further shows that the small molecules act on a very narrow site of RBD. Our data paves a way to design strain specific small molecule drugs as antagonists for Omicron as treatment modality for this variant of concern.

## Notes

### Competing Interest Statement

The authors have declared no competing interest.

